# Closed-loop control and recalibration of place cells by optic flow

**DOI:** 10.1101/2022.06.12.495823

**Authors:** Manu S. Madhav, Ravikrishnan P. Jayakumar, Brian Li, Francesco Savelli, James J. Knierim, Noah J. Cowan

## Abstract

Understanding the interplay between sensory input, endogenous neural dynamics, and behavioral output is key toward understanding the principles of neural computation. Hippocampal place cells are an ideal system to investigate this closed-loop interaction, as they are influenced by both self-motion (idiothetic) signals and by external sensory landmarks as an animal navigates its environment^1–9^. To continuously update a position signal on an internal “cognitive map”, the hippocampal system integrates self-motion signals over time^10,11^. In the absence of stable, external landmarks, however, these spatial correlates of neuronal activity can quickly accumulate error and cause the internal representation of position or direction to drift relative to the external environment^1,5^. We have previously demonstrated that, in addition to their known roles in preventing and/or correcting path-integration error, external landmarks can be used as a putative teaching signal to recalibrate the gain of the path integration system^6^. However, it remains unclear whether idiothetic cues, such as optic flow, exert sufficient influence on the cognitive map to enable recalibration of path integration, or if instead an unambiguous allocentric frame of reference, anchored by polarizing landmark information, is essential for path integration recalibration. Here, we use principles of control theory^12,13^ to demonstrate systematic control of place fields by pure optic flow information in freely moving animals by using a neurally closed-loop virtual reality system that adjusts optic flow speed as a function of real-time decoding of the hippocampal spatial map. Using this “cognitive clamp”, we show that we can not only bring the updating of the map under control of the optic flow cues but we can also elicit recalibration of path integration. This finding demonstrates that the brain continuously rebalances the influence of conflicting idiothetic cues to fine-tune the neural dynamics of path integration, and that this recalibration process does not require a top-down, unambiguous position signal from landmarks.

## Main

The spatial firing fields of hippocampal place cells are determined by allothetic inputs (such as visual landmarks and environmental boundaries) and path integration of idiothetic inputs (such as optic flow and vestibular signals)^9^. Decades of research have provided detailed insight into how allothetic cues can exert prercise control over the firing of place cells^3,6,7,14–17^. However, much less is understood about the mechanisms by which idiothetic cues affect place cells because, in the absence of landmarks, the updating of the map is unstable—when only idiothetic cues are available, the internal representation drifts and rapidly becomes unbound to the world frame of reference^1,2,11^.

Control theory provides a basis for stabilizing unstable systems and thus provides a powerful experimental arsenal to disentangle the elements of neural computation^12,13,18–22^. Famously, the voltage clamp allowed Hodgkin and Huxley to stabilize the membrane potential at a constant reference in order to pinpoint the roles of individual ion channels^23^. More recently, a growing body of literature has garnered new insights into neural computation by using control engineering to close feedback loops on neural representations^24–27^. Here, we extend the application of control theory to high order spatial representations; specifically, we introduce a “cognitive clamp” that maintains at a desired reference an essential cognitive variable for forming the hippocampal cognitive map, the gain of the path integrator. This gain relates self-motion information from idiothetic cues to an updating of position on the internal hippocampal representation.

We used a unique, immersive planetarium-style virtual reality (VR) apparatus (the “Dome”^28^; Fig. 1a) to provide pure optic flow input to a running rat while recording hippocampal place cells. Our study consisted of two conditions. In *open-loop* experiments, the optic flow cues in the Dome were tied to the movement of the rat such that its velocity with respect to the cues could be directly controlled via a pre-determined profile, building on human and other animal behavioral^29–36^ and neurophysiological^37–44^ experiments. In novel *closed-loop* experiments, the rat’s velocity relative to the cues was modulated in relation to neurophysiological activity in its hippocampus. This neural feedback was able to stabilize and recalibrate the path integrator gain in the absence of an absolute spatial reference frame typically provided by allothetic landmarks.

**Fig. 1.**
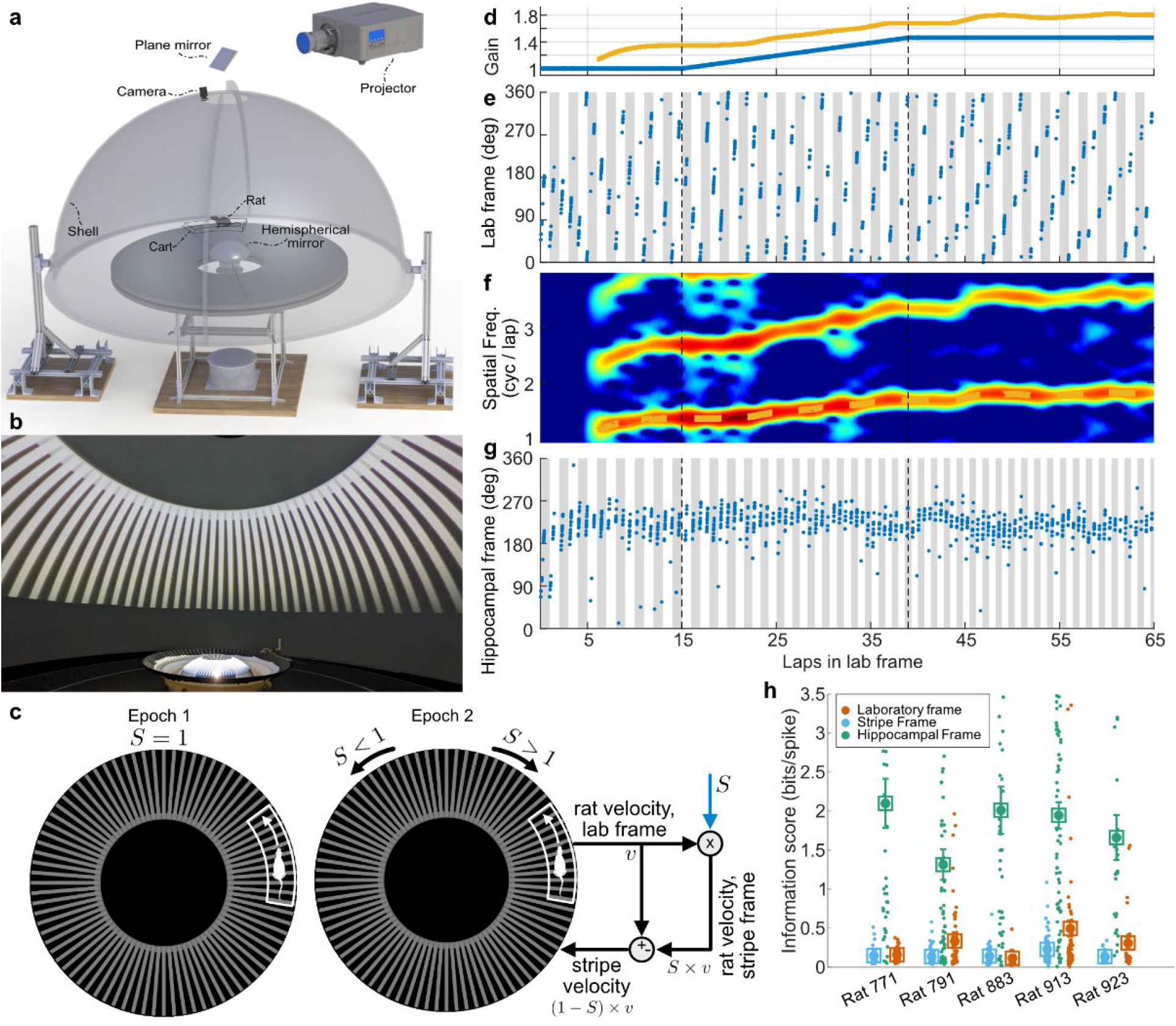
(a) VR Dome apparatus. Rats ran on a circular table surrounded by a hemispherical shell on which visual patterns were displayed by a projector, whose image reflected off a hemispherical mirror onto the inner surface of the shell. The rat ran within the bounds of an enclosure that was automatically moved to track the rat’s position as measured by an overhead camera. (b) Stripes projected within the dome. (c) The stripe gain, *S*, related the velocity of the rat with respect to the lab, *v*, and its velocity with respect to the stripe frames of reference, i.e. *S* × *v*. In Epoch 1, stripes were stationary (*S* = 1). In Epoch 2, stripes were moved in the same (*S* < 1) or opposite (*S* > 1) direction of the rat; specifically, the stripes were commanded to move at the difference (circle with +/-) between the rat’s velocity in the respective frames, namely *v* − *S* × *v* = (1*− S*) × *v*. (d-g) Hippocampal gain decoding. The *x* axis on all plots is the angular distance the rat ran on the table, in laps. (d) Stripe gain (*S*; blue) and hippocampal gain (*H*; yellow) during Epochs 1-2 in one session. *S* = 1 in Epoch 1 and was ramped up to and held at *S* = 1.46 during Epoch 2. *H* was decoded by our algorithm (see Supp. Fig 3). (e) Spikes from one unit (blue) plotted as a function of the rat’s angle *θ* (deg) on the table, relative to the laboratory frame of reference. Each gray or white vertical bar denotes 1 lap in the lab frame. The unit fired at the same location in the lab frame (*H* ≃ 1) for the first 2 laps but its place field began to drift backward on the track starting at lap 3 (*H* > 1). (f) Spatial spectrogram of firing rate of this unit computed with a 6-lap sliding window. The *y* axis denotes spatial frequency and color denotes power at each frequency at each spatial location. The dotted line shows the dominant spatial frequency tracked by our algorithm (i.e., *H*). The second harmonic of *H* is also evident. (g) Same spikes as (e) plotted in the hippocampal frame of reference (*Y* = ∫ *H dθ*, wrapped at 360°) with gray and white bars denoting laps in this frame of reference. Firing fields are aligned in the hippocampal frame of reference, indicating accurate decoding of *H*. (h) Spatial information scores in the hippocampal frame are significantly higher than those in laboratory and stripe frames for each rat (one session/rat with greatest number of units to avoid double counting, sessions with n= 43, 64, 51, 78, 28 units). Data are mean +/- s.e.m. with scores from individual units shown.

## Open-loop stripe manipulation influences hippocampal gain

Spatially alternating light and dark stripes were projected onto the inside of the Dome shell to provide a pure optic flow signal (Fig. 1b). There were no salient, polarizing landmarks in the Dome, and thus the rats were presumably forced to rely on idiothetic cues and path integration to maintain their sense of location as they ran laps. Rats typically ran counterclockwise (CCW) an initial 15 laps when the stripes were stationary (Epoch 1; Fig. 1d). After Epoch 1, we began rotating the visual stripes as a continuous, predetermined function of the rat’s movement through the environment. The stripes were moved according to a gain *S*, which determines the ratio of a rat’s speed relative to stripes to the rat’s speed in the lab frame. The stripes only moved when the rat moved. When *S* > 1, the stripes moved in the direction opposite to the rat’s movement; when *S* < 1, the stripes moved in the same direction as the rat; and when *S* = 1, the stripes did not move at all (Fig. 1c). For example, if a rat ran CCW, then with *S* = 2, the stripes moved at the same speed as the rat, but CW; with *S* = 0.5, the stripes moved at half the speed of the rat in the same CCW direction.

A typical open-loop session (N = 5 rats, 41 sessions, mean 22 units/session meeting place-field inclusion criteria; see Methods) is shown in Figs. 1d-g. For the first 15 laps, place fields drifted backward each lap. This drift is a consequence of the error that accumulates every lap when the animal must rely solely on path integration without any landmarks to prevent and/or correct drift. Importantly, this drift indicates that any landmarks in the environment (inside or outside the Dome) were insufficient for the rat to anchor its map. On lap 16, we began to rotate the stripes by ramping the stripe gain *S* up to a value of 1.461. The place fields drifted more swiftly, in the same direction as the stripe movement. This indicates that optic flow alone can qualitatively influence place fields^40^, similar to previous work with thalamic head direction cells in rats^39^ and retrosplenial spatial cells in mice^38^.

To quantify the drift of place fields over time, we estimated the gain of the hippocampus using an improved version of the population decoder in our prior study^6^. The hippocampal gain *H* can be thought of as the relationship between the animals’ physical movement through the world and the updating of its cognitive map. When *H* = 1, the firing pattern of a spatial cell repeats precisely once per lap; when *H* < 1, in contrast, the pattern repeats less frequently than once per lap, i.e. the rat’s position on its hippocampal map updates more slowly than the rat’s actual movement on the track (and vice versa when *H* > 1). The decoder works by determining the spatial frequency of place cells’ repeated firing fields as the animal traverses multiple laps (Fig. 1f; see Methods, Extended Data Fig. 3a). We found that the responses of place cells recorded in any given session were largely coherent during the manipulations of optic flow (Extended Data Fig. 3d). By integrating the decoded value of *H*, we can compute the position of the rat in its own internal reference frame, which we term the hippocampal frame. The rate maps of place fields calculated in the hippocampal frame (Fig. 1g) had greater spatial information than rate maps calculated in either the laboratory frame or the moving-stripe frame, whereas information in laboratory and stripe frames were not significantly different from each other (Fig. 1h; paired t-tests on mean information across rats: hippocampus vs. laboratory, t(4) = 8.50, p = 0.0010; hippocampus vs. stripes, t(4) = 11.94, p = 0.00028; laboratory vs. stripes, t(4) = -2.19, p = 0.094). These results indicated that the decoding of *H* was accurate and produced stable rate maps in the hippocampal frame of reference.

Pure optic flow cues exerted demonstrable influence over the hippocampal representation in all 5 rats (Fig. 2). Across sessions, we varied the stripe gain between 0.231 and 1.769 to produce a parametric description of how optic flow cues influenced the hippocampal spatial map. Fig. 2a shows a control session when *S* was maintained at 1 (blue line) (i.e., the stripes were stationary). The hippocampal gain *H* started out at approximately 1.09 in the first laps of Epoch 1, and this value gradually increased over the course of the session. Because *H* > 1, the place fields drifted on the track throughout the session (5 place fields recorded simultaneously in Fig. 2b), and the rate of drift slightly increased (i.e., the slopes of each cell’s lap-by-lap field location became steeper with increasing laps). When plotted in the hippocampal frame (Fig. 2b, bottom), each cell’s place field was stable, demonstrating that the hippocampal map drifted as a coherent unit. Fig. 2c,d shows an example session where *S*_final_ = 1.462. Here, *H* settled to a relatively steady value of *H*_baseline_ = 1.33 in Epoch 1, and then began to rise sharply in Epoch 2, paralleling the rapid increase in *S*. When *S* reached its final value (Epoch 3), *H* also stabilized, albeit at a higher value (*H*_final_ = 1.80), maintaining approximately its initial baseline offset. Fig. 2e,f shows a final example session in which *S* was decreased below 1. As was typical, *H* was greater than 1 in Epoch 1; when *S* was ramped down to 0.231 in Epoch 2, *H* decreased accordingly for the first ∼10 laps of Epoch 2. After that time, however, *H* appeared to reach a value of ∼1.06 in Epoch 2a, and then further decreased to *H*_final_ = 0.84 in Epoch 2b. Thus, the change in *H* followed the direction of the change in *S* but did not decrease below ∼0.84, even though *S*_final_ was a much lower value.

**Fig. 2.**
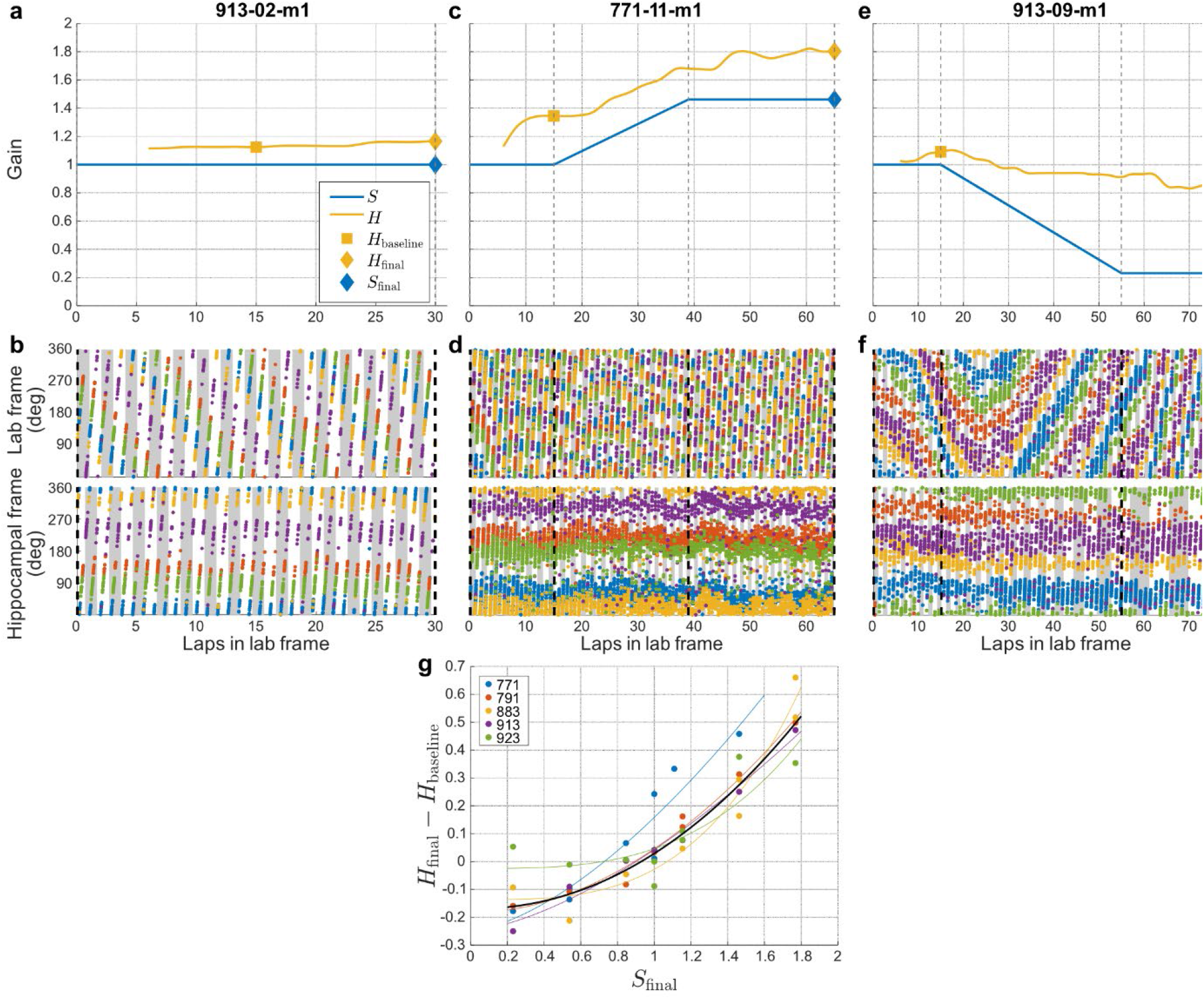
Effect of stripe manipulation on hippocampal place cells. (a) Gain curves for a session with *S* = 1 (stationary stripes; blue line). The hippocampal gain *H* (yellow line) drifted slightly, remaining above 1. No value for *H* is plotted for the first 6 laps because *H* is a 6-window average. (b) Spikes from five units (distinct colors) plotted in the laboratory frame (top) and hippocampal frame (bottom) for the session depicted in (a). All units are coherent with each other and drift at the same rate. They have stable firing fields (i.e., the fields are aligned horizontally) in the hippocampal frame. The alternating gray and white bars indicate laps in the respective frames of reference. (c) Gain curves for a session when *S*_final_ > 1. *H* initially increased and settled to a constant value towards the end of Epoch 1 with *S* = 1, but was driven upward when *S* was increased. (d) Same as (b), but for the session in (c). (e) Gain curves for a session when *S*_final_ < 1. *H* was driven downward when *S* decreased, but not to the same extent as upward manipulation. (f) Same as (b), but for the session in (e). (g) The change of hippocampal gain from its Epoch 1 baseline value is a nonlinear function of the stripe manipulation, parameterized by the final stripe gain *S*_final_. The relationship is captured by a power law 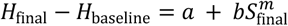 (fit parameters ± 95% CI: *a* = −0.17 ± 0.07, *b* = 0.20 ± 0.08, *m* = 2.12 ± 0.61; adjusted *r*^2^ = 0.88, n = 41, df= 38).

Across animals and sessions, the hippocampal gain at the end of Epoch 2 (after baseline subtraction; i.e., *H*_final_ − *H*_baseline_,) was strongly related to the final stripe gain *S*_final_ in Epoch 2 (Fig. 2g). However, as demonstrated in the examples in Figures 2a-f, the relationship was not linear. For *S*_final_ > 1, the relationship was approximately linear with a slope of 0.58 (*p* = 1.6 × 10^−5^, *n* = 16), showing that there was reliable, but incomplete, control of the hippocampal gain by optic flow cues. However, for *S*_final_ < 1, the slope was much less than for *S*_final_ > 1. A power law, 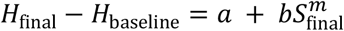, empirically provided a good fit to the data (Fig. 2g): (adjusted *r*^2^ = 0.88, *n* = 41, *df* = 38). This nonlinear relationship indicates an important asymmetry in the affordance of optic flow over the hippocampal gain in upward versus downward directions and is consistent with an asymmetric influence of optic flow reported in associated regions^36,38^.

## Closed-loop cognitive clamp stabilizes path integration

Despite the clear, bidirectional influence of optic flow on place cells, the precision of its control over the place fields was variable and offset by significant shifts of the baseline gain from session to session (Extended Data Fig. 4). This imprecision contrasts with the powerful control typically exerted by salient landmarks in the environment^3,9,14^, which was central to our prior demonstration that persistent conflicts with polarizing landmarks caused recalibration of the path integrator^6^. We investigated whether we could mimic the strong control by landmarks with pure optic flow information by using concepts from control theory to clamp the hippocampal gain to a desired value. Specifically, we created a neural feedback control loop in which CA1 place cell activity was used to adjust the experimental stripe gain, *S*, in real time to drive the hippocampal gain to an experimentally chosen desired value, *H*_desired_ (Fig. 3a).

**Fig. 3.**
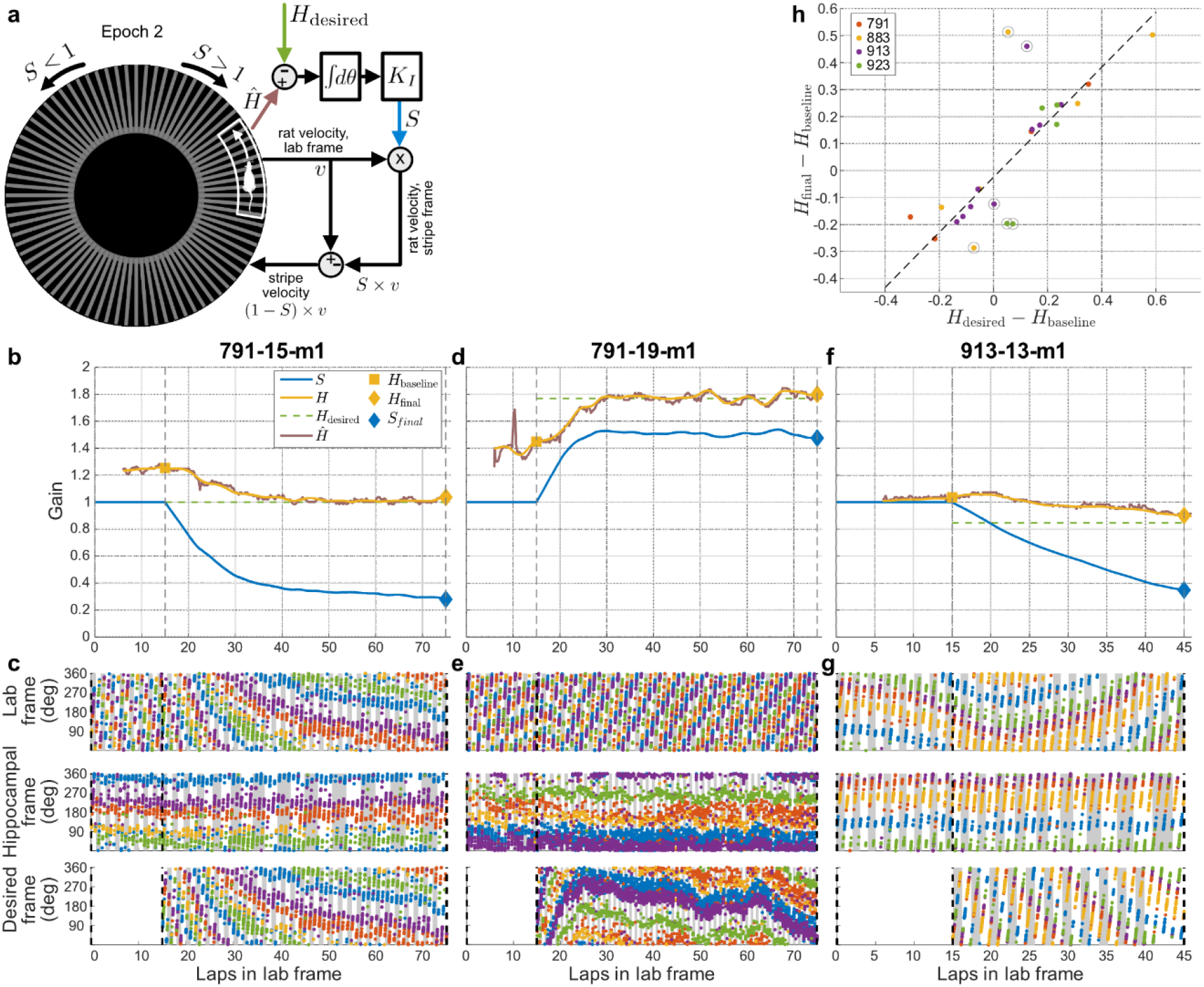
Cognitive clamp of hippocampal gain via optic flow. (a) Schematic of closed-loop controller. The calculation of the stripe velocity was the same as in Fig. 1(c), except that during Epoch 2, the stripe gain *S* was continually updated via an integral controller (see text); the controller was designed to clamp an estimate of the hippocampal gain 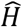 (see Extended Data Fig. 3) to the desired value *H*_desired_. (b) Example of closed-loop control to *H*_desired_ = 1. The stripe gain *S* (blue), real- time hippocampal gain estimate 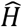 (brown), offline hippocampal gain *H* decoded after the experiment (yellow), and desired hippocampal gain *H*_desired_ (dashed green line) are plotted as a function of cumulative angular displacement in laps relative to the laboratory frame of reference. *H*_baseline_ denotes the time point used to measure the hippocampal gain prior to the onset of the closed-loop controller and *H*_final_ denotes the final hippocampal gain at the end of Epoch 2. In this session, the hippocampal gain in Epoch 1 (before lap 15) was ∼1.2. When the controller was activated, the stripe gain became increasingly lower, driving the hippocampal gain to *H*_desired_ = 1 by around lap 40 and maintaining it there throughout the remainder of Epoch 2. (c) Raster plots for 5 place cells for the session in (b). The top, middle, and bottom graphs show the place fields in the lab, hippocampal, and desired reference frames, respectively. The desired reference frame was computed by integrating the desired, constant hippocampal gain: ∫ *H*_desired_ *dθ* = *H*_desired_ × *θ* (since *H*_desired_ was constant). No points are shown for Epoch 1 in the desired frame as the controller was not activated until Epoch 2. Because *H*_desired_ = 1, the top (lab frame) and bottom (desired frame) graphs are identical during Epoch 2. (d) Example of closed- loop control successfully driving the hippocampal gain toward *H*_desired_ = 1.77 in Epoch 2 and maintaining it at this level for the remainder of the Epoch. (e) Raster plots for 5 place fields from experiment in (d). (f) Example of closed-loop control to *H*_desired_ = 0.85. In this example, the gain in Epoch 1 was only slightly higher than 1 and rising. When the controller was turned on, the rise in *H* was reversed and gradually moved closer to *H*_desired_. Although the controller was unable to bring *H* to *H*_desired_ by the end of the Epoch, *H* was nonetheless driven below 1 and still decreasing at the end of Epoch 2. (g) Raster plots for 5 place fields from experiment in (f). (h) *H*_final_ vs. *H*_desired_ for all 4 rats in the closed-loop control experiment. *H*_baseline_ was subtracted from both *H*_final_ and *H*_desired_. The strong linear fit, close to the unity line (dashed diagonal), demonstrates that in most sessions (especially for *H*_desired_ − *H*_baseline_ > 0), we were able to control the hippocampal gain strongly by optic flow cues alone via the cognitive clamp. Circled points denote uncontrolled sessions, where the controller was unable to bring *H* to *H*_desired_ (see Supp. Fig. 2). Note that7the linear fit is to all points.

This control scheme compares a real-time, neurally decoded estimate of the hippocampal gain, 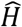, with the desired hippocampal gain, *H*_desired_, and feeds their difference back through an integral control law that automatically adjusts the stripe gain:

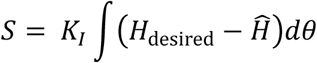

Here, *θ* denotes the cumulative, unwrapped angular displacement of the rat (measured in units of laps). The value of *S* was initialized to 1 at the beginning of Epoch 2. The controller constant *K*_*I*_, known as the integral gain in control theory, was designed to create a stable closed-loop system based on a simplified model that takes into account the 6-lap windowing of our real-time hippocampal gain estimate (see Methods). Intuitively, an integral control law continuously increases or decreases the strength of the control signal (i.e., *S*) until the feedback error is extinguished. The integral control law also created smooth changes in the stripe gain; that is, a gradual “ramp” emerged that is qualitatively similar to the pre- programmed stripe gain ramp presented in Epoch 2a of our open-loop experiments (see Fig. 2). This gradual ramp avoided sudden changes in optic flow velocity as might result from other control schemes (e.g., proportional or derivative controllers).

Our controller modulated *S* at 1-s intervals with *K*_*I*_ = 0.2 (see Methods). The control law was implemented on four animals across a total of 25 closed-loop sessions (mean 32 units/session). Fig. 3b depicts an example session in which *H* was initially greater than 1 (*H*_baseline_ = 1.253). The controller gradually reduced the stripe gain based on the integral control law, evidently causing a percept to the animal that it was moving progressively slower, until ultimately the hippocampal gain returned to unity (*H*_final_ = 1.037). In steady state, a population of simultaneously recorded place cells largely stabilized itself relative to the track (Fig. 3c), even in the absence of salient landmarks.

Our controller was successful for a large fraction of sessions in which *H*_desired_ > 1 (see Extended Data Fig. 2). Fig. 3d depicts an example in which the hippocampal gain was gradually ramped up to a desired value of 1.769 via the integral controller and stabilized around that value for approximately 45 laps (*H*_final_ = 1.800). As can be seen in Fig. 3e, a population of simultaneously recorded neurons became relatively stable in an artificial reference frame that rotated according to the desired reference frame, demonstrating the effectiveness of the control law. The control law was generally not successful in completely stabilizing to *H*_desired_ < 1, although there was often still an influence of the control law (Fig. 3f,g). This result parallels the relatively modest effect for *S* < 1 described earlier in open-loop experiments (see Fig. 2e). To assess the overall control law’s effectiveness, we correlated *H*_final_ at the end of Epoch 2 with *H*_desired_, after subtracting baseline from both variables (Fig. 3h). There was a linear relationship close to unity between these values, demonstrating that our neurally closed-loop controller can was able to systematically command the rate of updating of the hippocampal map using purely optic flow cues (slope = 1.02, *r*^2^ = 0.61, p = 3.76 × 10^−6^).

## Recalibration of path integration without landmarks

Previously, we showed that imposing a sustained conflict between idiothetic path integration and movement relative to *allothetic* cues (i.e., landmarks) induced recalibration of the path integration gain^6^. Here, we leveraged our cognitive clamp to investigate whether such recalibration relied on polarizing landmark information, or if the hippocampal network would also recalibrate in the face of sustained conflicts between distinct *idiothetic* cues. Such an effect would demonstrate a previously unknown degree of plasticity in how various idiothetic sources either mutually calibrate each other or recalibrate a downstream circuit, to continuously fine-tune the path integrator without polarizing landmarks.

As described above, we used our closed-loop controller to induce a conflict between optic flow and other idiothetic cues that were not manipulated (e.g., vestibular, motor copy, proprioception) (Epoch 2). To test for optic flow-based recalibration, we next extinguished the stripes (Epoch 3) and examined the hippocampal gain, *H*. We restricted our analysis to cases where the control law was successful in driving *H* to the desired value (see Methods and Supp Fig. 2; 10 strongly controlled and 8 modestly controlled sessions). One rat (#923) was eliminated from further analysis as no sessions for that animal were strongly controlled. Two example sessions (Fig. 4a, strong control and Fig. 4b, modest control) illustrate the effect of the recalibration, as there is a residual effect in Epoch 3 of the gain control manipulation carried out in Epoch 2. This effect was observed across sessions for all three animals, as there was a strong, linear relationship between the desired hippocampal gain and the measured hippocampal gain after the stripes were extinguished (Fig. 4c); note that the statistical tests were performed after subtracting the baseline offset from each session, a measure that ensures that the experimentally prescribed *change* in gain (relative to baseline) was correlated with a *change* in the hippocampal gain after stripes were extinguished.

**Fig. 4.**
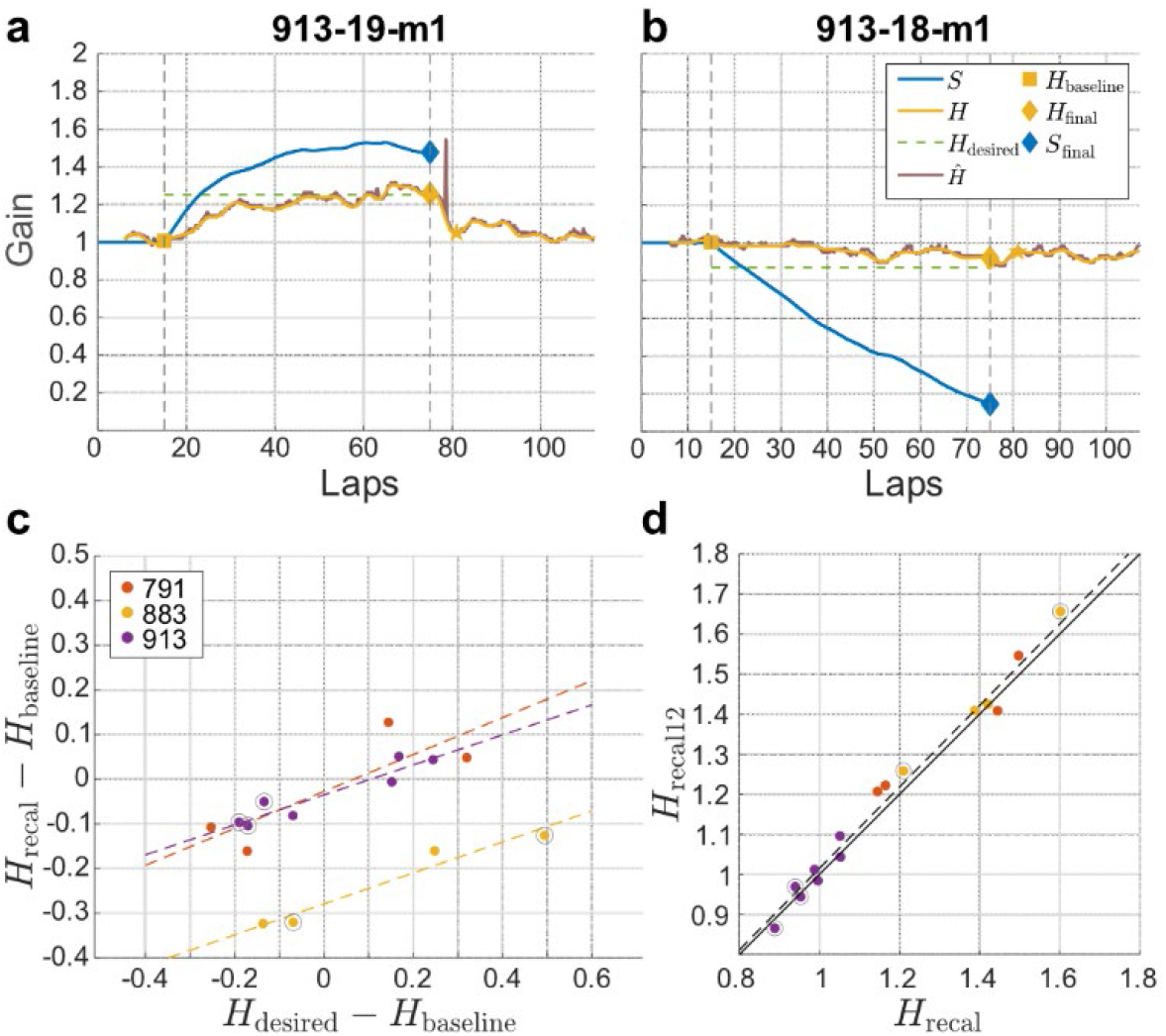
Recalibration of path integration gain via the cognitive clamp. (a) Example of recalibration to a gain *H* > 1. The closed-loop controller drove the hippocampal gain *H* to *H*_desired_ = 1.25. When the stripes were turned off at lap 75, *H* was reduced to ∼1.09, which was higher than the baseline gain in Epoch 1 (*H*_baseline_ = 1.01). (b) Example of recalibration when *H*_desired_ < 1. This example comes from the same animal as (a) on the previous day, in which *H*_desired_ was set at 0.87. The hippocampal gain was maintained at *H* = 1 for 15 laps in the absence of landmarks, and the gain was reduced to slightly below 1 for ∼20 laps when the controller was activated. At lap 45, the hippocampal gain started to decrease further and continued to decrease toward *H*_desired_. When the stripes were turned off at lap 75, the hippocampal gain was maintained at this value. (c) Hippocampal gain after recalibration (*H*_recal_) as a function of the closed-loop controller desired gain (*H*_desired_) for the sessions from 3 rats in which the stripes strongly controlled the hippocampal gain in Epoch 2. For all 3 rats, there was a strong, linear relationship between *H*_recal_ and *H*_desired_ (with *H*_baseline_ subtracted from both variables). Modestly controlled sessions are circled in gray. For each rat, slopes = 0.41, 0.35, 0.34; *r*^2^ = 0.64, 0.94, 0.87; *n* = 4, 4, 7. (d) Stability of *H*_recal_ over laps. There was a strong relationship close to unity between the values of H_recal_ measured 6 laps and 12 laps after the stripes were turned off (slope = 1.02, r^2^ = 0.99, p = 3.98 × 10^−16^, n = 18).

We examined whether the recalibration effect was persistent by tracking the hippocampal gain across Epoch 3 (stripes off). There was a near one-to-one correspondence between the hippocampal gain at laps 6 and 12 in Epoch 3 (Fig. 4d, slope = 1.02, r^2^ = 0.99, p = 3.98 × 10^−16^, n = 18). This correspondence shows that the optic flow-based hippocampal gain manipulation induced a long-term recalibration of path integration with respect to the other idiothetic cues (e.g., vestibular cues, proprioceptive cues, or motor copy) that presumably drove the path integration process when the optic flow cues were severely diminished in Epoch 3.

## Discussion

We used a virtual reality apparatus with a freely moving rat to demonstrate systematic control of place cells by optic flow, an idiothetic cue that is often hypothesized to be a major influence on place cells but that has not been studied extensively and parametrically in this regard. Under natural conditions, salient landmarks and boundaries anchor the internal reference frame of the hippocampus, making it difficult to study the influence of idiothetic cues in isolation and almost impossible to quantify how conflicting idiothetic cues interact in updating the path integration computation^1,2,11^. The robust control of place fields with optic flow achieved with our cognitive clamp is analogous to the well-documented control exerted by allothetic cues such as landmarks^3,6,7,14–17^. Our previous work showed that such allothetic information can provide a teaching signal to recalibrate the path integration system^6^. In that case, the teaching signal—the landmarks—provided an absolute positional signal in its frame of reference that anchored the internal, hippocampal frame of reference. Idiothetic cues, in contrast, can only provide relative positional signals (i.e., updating a positional signal relative to the previous estimate of position). Does the path integration system require an absolute position teaching signal to calibrate its gain, or could relative signals from different idiothetic sources calibrate each other? A neurally closed-loop controller allowed us to establish that manipulation of optic flow can induce recalibration of the path integrator in a similar way to what we had previously shown by landmark manipulations. Indeed, in the subsequent absence of the controlling stripes, the hippocampal gain value was linearly related to the desired gain value to which the hippocampal reference frame was clamped.

Robust internal dynamics are a hallmark of the hippocampal circuitry. Our research shows that the internal dynamics of the path-integration network are constantly being fine-tuned in relation to potentially conflicting streams of idiothetic information. Importantly, a global, top-down teaching signal that binds the hippocampal frame of reference to an absolute external frame of reference is not required for recalibration. Instead, the internal dynamics are the reference frame against which idiothetic inputs are compared, providing an externally ungrounded teaching signal. The algorithm for multimodal integration is reminiscent of clock synchronization and recalibration^45^. In the presence of a trusted master timekeeper (e.g., an atomic clock), drifting clocks are ‘latched’ onto it, and their rates of drift are corrected—much like visual landmarks anchor the spatial representation and induce path integrator recalibration. In the absence of this master clock, synchronization algorithms rely on a network of clocks synchronizing and calibrating each other—much like optic flow influencing (without anchoring) the spatial representation and nevertheless inducing recalibration.

By stabilizing the hippocampal representation in the absence of allothetic landmarks, the neural closed-loop controller we developed opens the door for studying idiothetic inputs to the hippocampus with a degree of control previously reserved for studies of allothetic inputs. The present study used an online decoder and controller to calculate and manipulate the hippocampal gain, but future work will likely be able to decode the hippocampal representation of the actual position of the animal in real time^46,47^ and control directly the location of the hippocampal “activity bump”^48^ based purely on idiothetic cues. Furthermore, the relative influence of different idiothetic inputs can be determined in ways analogous to classic voltage clamp studies. That is, one idiothetic input (e.g., vestibular) can be manipulated systematically, and the other (e.g., optic flow) can be adjusted to counter the manipulation and clamp the hippocampal representation. The magnitude of the controller input required to clamp the representation is a measure of the relative strength of the two cues’ influence on the updating of position on the hippocampal map, much like the current required to maintain the voltage clamp at a set value indicates the relative current flow through various ion channels^23^. Such neurally closed-loop experiments that regulate or stabilize internal variables can generalize to other fields of cognitive neuroscience in which high-order neural representations (e.g., evidence accumulation, motor intentions, or attention) are influenced by, but not necessarily bound to, external sensory input but are instead dynamically modulated by internal variables.

## Methods

### Subjects

Five Long–Evans rats (Envigo Harlan; 3 males [numbers 771, 791, and 883] and 2 females [numbers 913 and 923]) were housed individually on a 12:12 h light:dark cycle. There were no obvious differences in results between male and female rats, and the data are reported by rat as appropriate in the results. All training and experiments were conducted during the dark portion of the cycle. The rats were 5–8 months old and weighed 300–450 g at the time of surgery. All animal care and housing procedures complied with National Institutes of Health guidelines and followed protocols approved by the Institutional Animal Care and Use Committee at Johns Hopkins University.

### Dome apparatus

To present visual landmarks and optic flow cues to the rat, we used our custom planetarium-like virtual reality apparatus, the Dome (see ^28^, for details on the design and construction). Briefly, rats locomoted near the outer periphery of an annular table (152.4-cm outer diameter, 45.7-cm inner diameter) centred within the dome. The image from a projector (G7500UNL, Epson Inc., Nagano Japan) fitted with a long-throw lens (ELPLM11, Epson Inc., Nagano Japan) was reflected off a plane mirror (152 mm × 152 mm × 12.7 mm, First Surface Mirror LLC, OH, USA) and a hemispherical mirror (254 mm dia., 150 mm radius of curvature, 40/20 surface quality, 1/4-wave accuracy, protected aluminum coating, Cumberland Optical, MD, USA) mounted at the centre of the Dome. The resulting image covered the inside surface of the Dome shell, providing a projected view to the rat 360° in azimuth and almost 90° in elevation. A near-infrared camera (GS3-U3–41C6NIR-C, FLIR, OR, USA; 2048×2048 px, 45 fps) with a wide-field lens (NMV-6M1, 6mm, F1.8, Navitar, NY, USA) provided an overhead view of the experiment. The 3D position and orientation of the head of the rat were detected in real-time (45 fps) through single-camera tracking^49^ of a set of markers mounted on the rat’s neural recording implant. The central rotating pillar of the dome was mounted on bearings. An enclosure, attached to the central pillar via radial boom arms, covered a 45° region around the rat. The central pillar along with the enclosure were moved using a motor in response to the rat’s position, such that the rat was kept near the center of the enclosure. The enclosure was moved only when the rat moved forward (counterclockwise) and not when it went backwards (clockwise) – this encouraged continuous forward running during behavioural training. A micro-peristaltic pump (RP-Q1, Taskago Fluidics, Aichi Japan) on the central pillar dropped liquid reward (50% diluted Ensure®) through a feed tube routed to the front of the enclosure. A plastic spreader and paper towels were attached to a third radial boom arm mounted to the central pillar opposite from the enclosure. This cleaning arm wiped up or spread out the scent of urine and uneaten food, as well as pushed feces off the table, reducing the salience and stability of local olfactory cues. All the non-projected visual cues available to the rat were either circularly symmetric (non-polarizing) or moved along with the rat.

### Projected visual cues

During Epochs 1-2, a set of 80 equally spaced white stripes was projected into the dome to form the optic flow cue. The stripes were each 1.5° wide and 40° high, centered at 45° elevation. The spacing between the stripes was 360°/80 = 4.5°. The stripes were set to 50% brightness. Stripes were present in all except the last epoch in both open- and closed-loop experiments. A circular band (elevation: 65°, brightness 40%) was projected in all epochs to provide circularly symmetric illumination inside the dome. During the first 15 laps, before Epoch 1, a set of stationary landmarks—identical to those used in ^6^—were superimposed over the stripes, and both stripes and landmarks were also stationary. Because this overlay of landmarks and stripes did not reliably provide strong cue control, likely because of the lack of visual salience of the landmarks against the striped background, this pre-Epoch-1 landmark condition was excluded from further analysis.

### Training

Over 2–3 days, we familiarized the rats to human contact and trained to wear a body harness (Coulbourn Instruments). The rats were placed on a controlled feeding schedule to reduce their weights to approximately 80% of their ad libitum weight, whereupon they were trained to run for a food reward (50% diluted Ensure) on a training table in a different room from the experimental room. The training table had the same dimensions as the table inside the dome, but no enclosure or other automated systems. Reward droplets were manually placed at arbitrary locations on the track in the path of the running rat, and the experimenter attempted to lengthen the average interval between rewards to maintain behaviour while delaying satiation. Training continued until the rats consistently ran 40 laps without intervention or encouragement from the experimenters. Training usually took 2–3 weeks.

### Electrode implantation and adjustment

After training, rats underwent stereotactic surgery where they were implanted with hyperdrives containing 16 independently movable nichrome tetrodes, the tips of which were gold-plated to an impedance of ∼ 150 kΩ using a nanoZ electroplating system (White Matter LLC, Seattle WA USA). The hyperdrives were fabricated in the laboratory using an in-house design and used a 72-channel interface board (EIB-72-QC, Neuralynx, Bozeman MT USA). Following surgery, 30 mg of tetracycline and 0.15 ml of a 22.7% solution of the antibiotic enrofloxacin were administered orally to the rats each day. After at least four days of recovery, we began slowly advancing the tetrodes and resumed food restriction and training within seven days of surgery. Once the tetrodes were close to CA1, they were advanced less than 40 µm per day. The location of each tetrode relative to the CA1 pyramidal cell layer was judged using the polarity of sharp waves and intensity of ripples in the electroencephalogram (EEG) signal captured on one electrode of each tetrode, as per well-established procedures. Tetrodes were judged to be placed correctly when ripples were intense and multiple units were visible on the pairwise electrode projections of spike amplitudes.

### Post-surgery training

During days of electrode advancement, we simultaneously food-restricted the rats. On (typically) day 3 of food restriction, we placed them into the Dome while wearing the body harness. A magnetic pad attached to the harness was used to mount a 6.4-mm marker (Optitrack, Corvallis OR USA) to track the position of the rat and actuate the enclosure surrounding the rat in real time, so that the rat remained near the centre of the enclosure. To encourage movement in only one angular direction, the enclosure was never moved clockwise. Thus, as the rat approached the front (counterclockwise end) of the enclosure, it moved forward. When the rat approached the back (clockwise end) of the enclosure, it did not move, thereby blocking the path of the rat. A pump dropped liquid reward in front of the running rat, which prompted the rat to move forward and thus move the enclosure. Reward was dropped at a spatial interval picked from a uniform distribution around a mean interval that varied across days depending on the rat’s performance. Within a few days post-surgery, rats learned to run continuously and obtain food reward. The mean interval of reward was gradually increased to maintain running performance until the pre-surgery performance criterion was reached once again (typically 7-10 days).

### Neural recording

Once the tetrodes were judged to be in CA1 and the rat was again running at least 40 laps inside the dome, the experimental sessions began. During sessions, a unity-gain neural recording headstage (EIB-72-QC, Neuralynx, Bozeman MT USA) was attached to the implanted hyperdrive. The neural signals passed through the commutator and were filtered (600–6,000 Hz), digitized at 30 kHz, and recorded on a computer running the Neuralynx Cheetah 5 recording software. Simultaneously, EEG data from one channel of each tetrode was filtered (1–475 Hz), digitized at 30 kHz, and stored on the computer. Pulses sent from the experiment-control computer were time-stamped and recorded as events on the neural-recording computer to enable the post hoc synchronization of the data streams recorded on the two computers. Procedures for synchronizing and associating signals to the behavioural data are detailed in (Madhav et al., 2021). For experimental sessions, instead of the single large marker attached to the body harness, a set of smaller (3mm, 4 mm) markers were placed in a rigid arrangement around the recording headstage. This allowed our custom algorithm to track the 3D position and orientation of the constellation of markers with higher accuracy and robustness^49^. Thus, the rat did not need to wear the harness during sessions.

### Experimental control

Three computers were used to run the experiment. Their purposes were: (1) general experiment control, (2) neural recording, and (3) video tracking and neural recording. Multiple independent programs, called nodes, performed each of these tasks and communicated to a master node running on Computer #1 and to each other through a software framework called Robot Operating System (ROS)^50^. Details on the hardware and software integration and experimental control are available in ^28^.

### Real-time firing rate computation

A python ROS node on the neural recording computer used the NetCom Application Programming Interface (API) to receive real-time neural data, and ROS APIs to receive tracked rat positions. Occupancy of the rat and spike counts from each tetrode were collected into 5° spatial bins covering a region 6 laps prior to the current angular position of the rat (6 × 360/5 = 432 bins). Rat velocities were computed at 100 Hz, and a count was added to the current occupancy spatial bin if the velocity was above 5 deg/s (≈5 cm/s). Spikes from each tetrode were tested for high, correlated amplitudes (indicating noise) and then counted into their respective spatial bins if the current velocity was above threshold. Spike counts were divided by occupancy, and the resulting 432 firing rates (spikes/s) were made available at 1 Hz. These firing rates were used by the online spectral decoder described below, to decode hippocampal gain (Extended Data Fig. 3c) for each tetrode. Tetrodes that had no visible neurons or had noisy recordings were excluded by the experimenter using a manual interface, and the median of the gain estimates from the remaining tetrodes was termed the online hippocampal gain 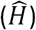 and used to manipulate stripe gain *S* during Epoch 2 of closed-loop sessions.

### Spectral Decoding

The algorithm for spectral decoding of hippocampal gain is detailed in Extended Data Fig. 3. The core algorithm remained the same as in our previous paper^6^, but updates (described below) were made to improve the robustness of gain estimates. Briefly, we used the spatial periodicity of firing rates of place fields on a circular track to compute the spatial frequency of the population representation. For a stable spatial representation in the laboratory frame, a typical CA1 place cell would exhibit one firing field that repeats every lap, hence the spatial frequency of firing is 1 cycle/lap. If a cell fired more (or less) than once per lap, its spatial frequency would be > 1 (or < 1) cycles/lap. The spatial frequency of firing is termed the hippocampal gain of each place cell.

In this version of the decoder, we improved the threshold mask used to enhance the signal-to-noise ratio of the spatial frequency content. The spatial spectrogram of the firing rate curve of each unit was first thresholded to the 80% percentile of its power in each spatial window. Contiguous regions above the percentile threshold were identified (MATLAB *regionprops* function). Noise regions tended to lack structure and agglomerated into punctate roundish blobs while the parts of the spectrogram denoting spatial frequency traces were larger in pixel count and more elongated. Given this, regions which were below a pixel area of 70000 and an aspect ratio of 17 were removed from the mask. This thresholding mask was then applied to the sharpened spectrogram.

There were instances when the power in the fundamental trace failed to exceed the threshold described previously, causing the maximum-energy trajectory to follow a harmonic instead of the fundamental. An assumption was made that if the gain of a cluster at a particular spatial window is a harmonic of another cluster, the two clusters were likely from the same periodic signal. Harmonics were identified by taking the pairwise ratios between the gain estimates of all clusters at each spatial window. Ratios that were close to integer values indicated that the numerator is likely a harmonic of the denominator and were divided by this integer. These harmonic-corrected gain estimates from the individual clusters were binned in the space of unwrapped angular position and spatial frequency to identify if the set of gain estimates had a coherent grouping (all cluster gain estimates fell within a 0.05 mean absolute error of each other) or if multiple subpopulations existed.

#### Offline decoding

The spectral decoder was run with a spatial window on each sorted unit passing inclusion criteria (see Data Analysis, below). For each 5° spatial bin, the firing rate for each unit was calculated by dividing the number of spikes fired by that unit by the amount of time the rat spent in that bin when it was moving > 5°/s. For each unit and for each bin, the hippocampal gain was the spatial frequency estimated by the spectral decoder on the 432 firing rates corresponding to the 6-lap window prior to that bin. The population hippocampal gain *H* for each bin was computed as the median of these estimates across units.

#### Online decoding

During online decoding, the spectral decoder was run at 1 Hz on the 432 spatially binned firing rates for each tetrode corresponding to the 6 laps prior to the rat’s current angular location. Thus, every second, a hippocampal gain estimate was generated for each tetrode. The population online hippocampal gain estimate 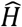 was the median of these estimates across the tetrodes chosen by the experimenter.

### Closed-loop controller design

We hypothesized that we would be able to control the hippocampal gain *H* to a desired value by manipulating the stripe gain *S* during an experimental session, in response to an online decoded gain 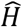. Open-loop experiments demonstrated that changes in *S* produced changes in hippocampal gain *H*, but that this change effect is nonlinear. From Fig. 2g, we observed that, to a reasonable approximation

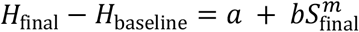

Without stripe manipulation (*S*_final_ = 1), we do not expect *H*_final_ to deviate significantly from *H*_baseline_. This is also supported by the data (Paired t-test, *H*_final_ vs. *H*_baseline_ in open-loop sessions where *S*_final_ = 1: *t*(6) = −0.90, *p* = 0.40). Thus, for the purpose of developing a feedback controller, we make the simplifying assumption that *a* = −*b*. We further assume a quadratic relationship (*m* = 2). These assertions are supported by curve fits across rats in Fig. 2g; (Paired t-test, *a* vs. −*b*: *t*(4) = 1.78, *p* = 0.15; *m* vs. 2: *t*(4) = 0.79, *p* = 0.048). (Our failure to reject the null hypotheses in these low-sample datasets is only used to furnish a simplified model that is sufficiently expressive for neurally closed-loop control design, not make definitive conclusions about the nature of the parameters.) Under these assumptions, we get the simplified model:

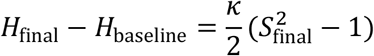

Where 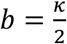. This corresponds to the integral equation:

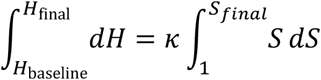

Ignoring constants of integration, the model relating the change in stripe gain *dS* to the hippocampal gain *dH* is thus

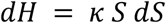

At the initial value of *S* = 1, the linearized system dynamics is given by *dH* = *k dS*. We designed a controller for this simplified system that reduces the error between the decoded gain 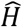 and the desired value *H*_desired_. In scenarios where a control input (*S*) is used to drive a system state (*H*) to a desired state *H*_desired_, a proportional controller can be used. For such a controller, the input is proportional to the error between 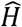 and *H*_desired_.

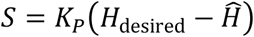

In our case, at a particular 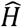 and *H*_desired_, the error would remain constant, thus *S* will be constant. Thus, *dS* = 0 which means that *dH* = 0 as well. Moreover, proportional control can require large gains to reduce the error, and large gains would lead to rapid changes in stripe movement such that the virtual environment may no longer have appeared to be stable to the animal. Due to these reasons, we chose an integral controller, known to be able to eliminate steady-state errors under appropriate conditions:

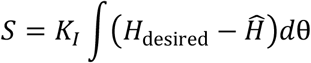

Here, the integration is initiated at *S*_0_ (*S*_0_ = 1 in our case, the value of the stripe gain at the beginning of Epoch 2). The term *K*_*I*_ is the “integral gain” (terminology from control theory, not to be confused with other ‘gains’ in the manuscript), and *d* is the angle of the rat on the table.

The block diagram of the feedback system consisting of the rat and controller is shown in Extended Data Fig. 7. The feedback loop consists of the controlled “plant” *P* (in this case, the hippocampal circuit), the integral controller *C* and the feedback, which is our decoder, represented by a moving average over a window of 6 laps. We performed a Nyquist stability analysis to determine the range of integral gain *K*_*I*_ over which the controller would be stable. When the frequency of the input signal *s* = *j*ω is swept from 0 to ∞, if the loop gain *L* intersects the real axis at a point less than -1, the system is unstable. To determine this point:

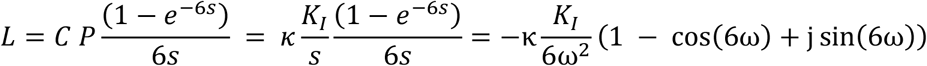

Here, 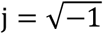. Setting Im[*L*] = 0, the first point of intersection with the real axis is 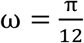.

The intersection with the real axis is 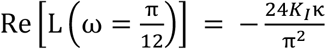.

To maintain stability, this value needs to be less than -1, thus it requires 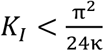

According to our fit in Fig. 2g, *b* = 0.2, and thus *k* = 0.4; hence the condition for stability is *K*_*I*_ < 1.03. We used *K*_*I*_ = 0.2 in our closed-loop sessions, staying well within this margin of stability.

### Stripe gain selection and ramp rates

In open-loop sessions, rats ran 15 laps with stripes on and stationary (Epoch 1, Figs. 1-2). In Epoch 2, *S* was increased or decreased to *S*_final_. The values of *S*_final_ were chosen to be of the form, 1 ± *n*/13 with *n* = 2, 6, 10, resulting in gains of 0.231, 0.539, 0.846, 1.154, 1.462 and 1.769. These values with a prime denominator were chosen to reduce ambiguity between frames and ensured that during Epoch 3 the position of the rat relative to the laboratory and stripe frames of reference aligned only once every 13 laps. Gains were changed at a constant rate of 1/52 per lap, such that the length of Epoch 2 was 8, 24 and 40 laps for n = 2, 6 and 10, respectively. The sessions were not randomized; the gain for each session was selected such that gains were rarely repeated in consecutive sessions, and the gain manipulation typically increased in magnitude over consecutive sessions for any given animal. The investigators were not blinded to allocation during experiments and outcome assessment. No statistical methods were used to predetermine sample size.

In closed-loop sessions, Epoch 1 was identical to open-loop sessions. The hippocampal gain decoder was initialized and the gain estimate 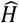 was monitored during Epoch 1, but not used for cue manipulation. *H*_desired_ was either specified before the session, in which case it was chosen from the aforementioned values of the form 1 ± *n*/13, or it was set to be a constant offset of ±0.25 or ±0.5 from the estimated value of *H*_baseline_. In either case, the *H*_desired_ value was set and not modified once the controller was initialized at the beginning of Epoch 2.

### Data analysis

#### Spike sorting

For each triggered spike waveform, features such as peak, valley and energy were used to sort spikes using a custom software program (WinClust; J.J.K.). Cluster boundaries were drawn manually on two-dimensional projections of these features from two different electrodes of a tetrode. We mostly used maximum peak and energy as features of choice; however, other features were used when they were required to isolate clusters from one another. Clusters were assigned isolation quality scores ranging from 1 (very well isolated) to 5 (poorly isolated), agnostic to their spatial-firing properties. Only clusters rated 1–4 were used for quantitative analyses including offline estimation of hippocampal gain.

#### Inclusion criteria

To be included in the quantitative analyses, sessions were required to meet the following criteria: sessions with stripe manipulation must have been run all the way to completion, i.e. the rat finished the session (Epoch 1-3, and removed after stripes were extinguished), and there were no major behavioural issues or long manual interventions during the session, as per our experimental notes. Session inclusion was determined before performing any of the statistical analyses across sessions detailed in this manuscript. For the 66 sessions that met these criteria, spikes that occurred when the movement speed of the rat was less than 5°/s (about 5 cm/s) were removed.

#### Closed-loop control categorization

Four authors (MM, RJ, JK, NC) categorized closed-loop control based on careful evaluation and discussion of each session’s hippocampal gain trajectories, blind to the outcome—recalibration or not—of the individual sessions. The sessions were placed in one of three categories: strong control, modest control, and uncontrolled (see Supp Fig. 2). Only strongly and modestly controlled sessions were included for recalibration analysis.

### Histology

Once experimental sessions were complete, rats were transcardially perfused with 3.7% formalin. The brain was extracted and stored in 30% sucrose-formalin solution until fully submerged. For 4 rats, the brain was sectioned coronally at 40 µm intervals. The sections were mounted and stained with 0.1% Cresyl violet, and each section was photographed. These images were used to identify tetrode tracks, on the basis of the known tetrode bundle configuration. A depth reconstruction of the tetrode track was carried out for each recording session to identify the specific areas in which the units were recorded. For one rat, we optically cleared the whole brain using the AdipoClear+ protocol^51^. The cleared brain was imaged using a lightsheet microscope (Ultramicroscope, LaVision BioTec, Bielefeld Germany) and the tetrode tracks were visualized in the autofluorescence channel using Imaris software to identify areas where units were recorded.

### Statistics

Parametric tests were used to determine statistical significance. Pearson product-moment correlations were used to test the linear relationship between variables. For non-linear relationships, data was fit using a nonlinear least-squares (MATLAB *fit* function) with specified model structures, and goodness-of-fit statistics (residual degrees of freedom, *df* = *n* − *m*, where *n* is number of data points and *m* is number of parameters, as well as *df*-adjusted coefficient of determination (*R*^2^) are reported). Paired, two-sided t-tests were used to compare information scores in laboratory, stripe and hippocampal frames of reference, which assumes normality. To prevent sampling the same cells across days for this analysis, the experimental session with the greatest number of units was chosen for each rat and for each tetrode. Mean and Standard Error of the Mean (s.e.m.) were used to plot information scores across units.

## Code availability

Custom code was written to analyse the datasets used in this study, and to generate figures for this manuscript. This codebase is versioned, and uses several third-party packages, the license files for which are included with the respective code. Access to the codebase can be provided by the corresponding author.

## Reporting summary

Further information on research design is available in the Nature Research Reporting Summary linked to this paper.

## Data availability

The datasets used in this study are available from the corresponding author upon reasonable request.

## Acknowledgements

We thank Eric Fortune for providing valuable comments on the manuscript; Kelly Wright, Kimberly Nnah, Audrey Branch, Marissa Ferreyros, Shahin Lashkari, Bharath Krishnan, Balazs Vagvolgyi, and Vyash Puliyadi for technical assistance; and H. Tad Blair for useful discussions. This work was supported by U.S. Public Health Service grant R01 NS102537 (N.J.C., J.J.K., F.S.) from the NINDS, by the Johns Hopkins University Kavli Neuroscience Discovery Institute (M.S.M.), and by a Discovery Award from the Johns Hopkins University (N.J.C, J.J.K.).

## Author Contributions

M.S.M., R.P.J., F.S., J.J.K, and N.J.C. conceived and designed the study. J.J.K. and N.J.C. supervised all aspects of the experiments and analysis. R.P.J. and M.S.M. designed and constructed the apparatus, performed experiments, and analyzed the data. B. L. performed experiments and analyzed data. M.S.M., R.P.J., B.L., J.J.K. and N.J.C. wrote the paper and F.S. provided critical feedback.

---

**Extended Data Fig. 1.**
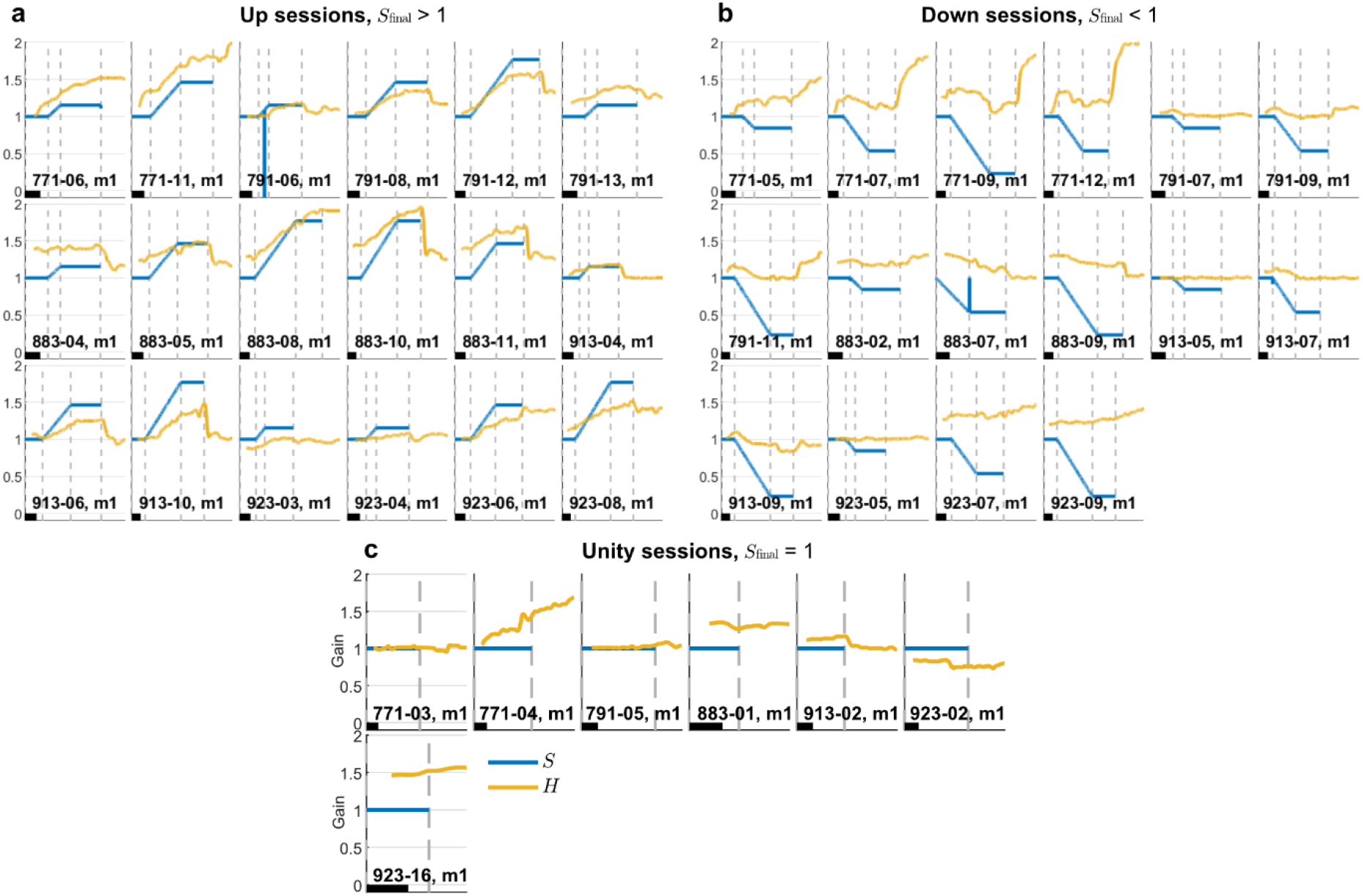
Gain dynamics during all open-loop sessions. Each plot represents a single session (titled as “Rat-Day, Session”, 41 sessions across N=5 rats). X axis is number of laps the rat ran on the table and Y axis is gain. The black scale bar in each plot denotes 10 laps. Applied stripe gain (*S*; blue) plotted with decoded hippocampal gain (H; yellow). Sessions are grouped by the final stripe gain: (a) Up sessions (*S*_final_ > 1), (b) Down sessions (*S*_final_ < 1), and (c) Unity sessions (*S*_final_ = 1). Dashed vertical lines indicate boundaries between Epochs 1, 2a, 2b and 3 for Up and Down sessions, and between Epochs 1 and 3 for Unity sessions. Blips in the *S* curve in sessions 791-06, m1 and 883-07, m1 were the result of momentary software errors.

**Extended Data Fig. 2.**
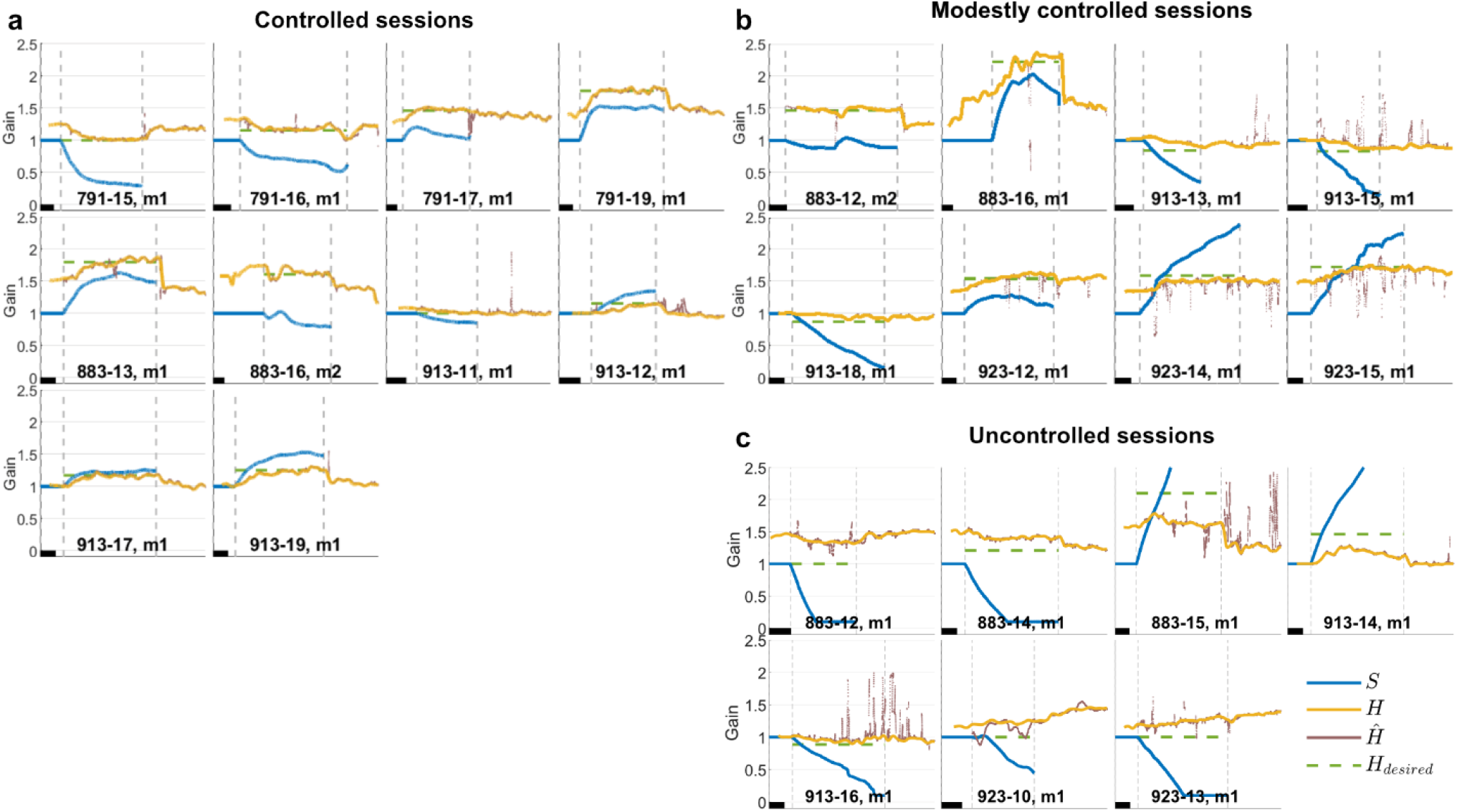
Gain dynamics during all closed-loop sessions. Each plot represents a single session (titled as “Rat-Day, Session”, 25 sessions across N=5 rats). X axis is laps the rat ran on the table and Y axis is gain. The black scale bar in each plot denotes 10 laps. Applied stripe gain (*S*; blue) plotted with offline-decoded hippocampal gain (*H*; yellow) and hippocampal gain estimated online using unsorted spikes (*H*; brown). This estimated value was driven to a constant desired value during the session (*H*_desired_; green dashed line). Dashed vertical lines indicate boundaries between Epochs 1, 2 and 3. Data is sorted into three groups based on manual inspection of our success in achieving consistent closed-loop control, i.e., how closely did the yellow line (*H*) match the dashed green line (*H*_desired_)?: (a) strongly controlled sessions, (b) modestly controlled sessions, and (c) uncontrolled sessions. Note that 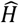 is a real-time estimate that depended on neural noise inherent in multi-unit electrophysiology and varied in quality day-to-day. 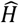 was utilized only in Epoch 2 and even then, our slow-moving integral controller mitigated the effects of momentary noise in the estimate.

**Extended Data Fig. 3.**
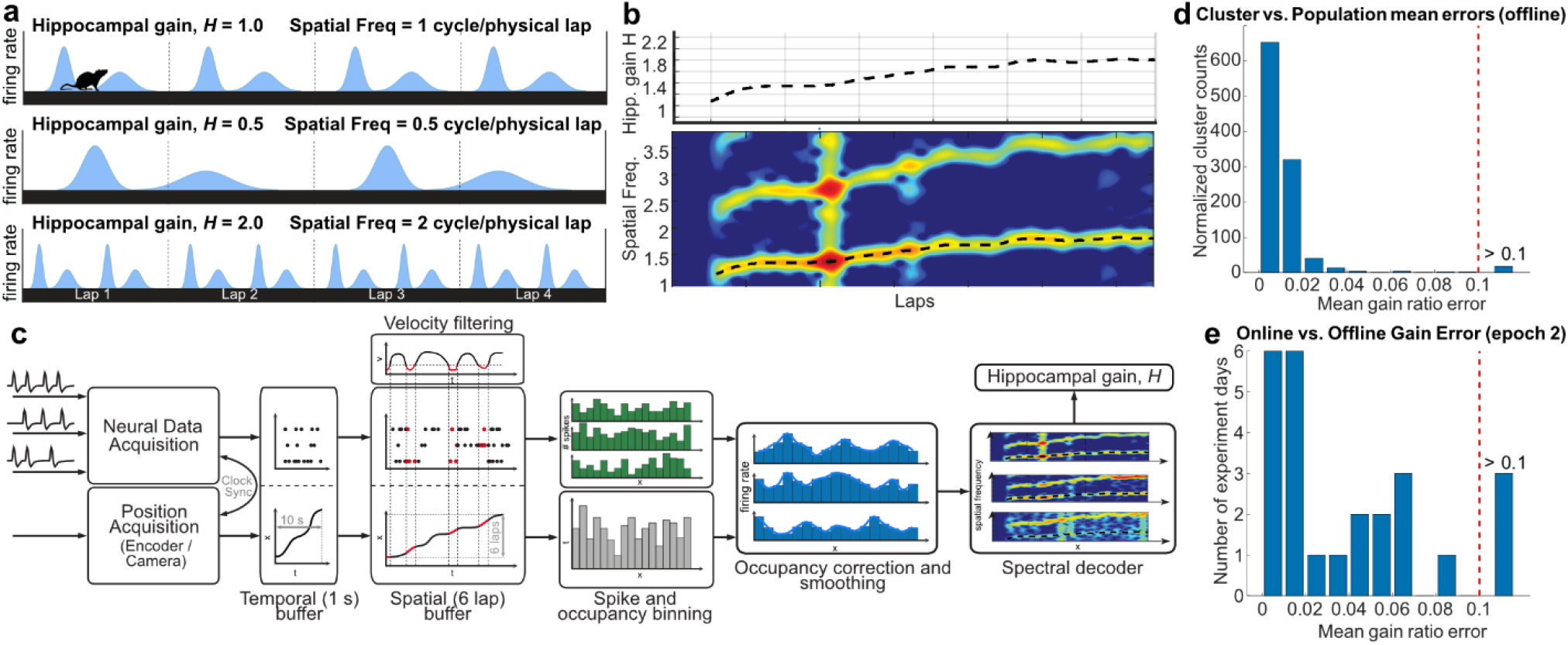
Hippocampal gain decoding. (a) If a spatially tuned cell has a characteristic spatial tuning that repeats once per physical lap, the firing rate of the cell exhibits a spatial frequency of *H = 1* per lap. Illustration of the firing of a spatially tuned cell for three values of hippocampal gain, *H*. (b) Reproduction of Figure 1d,f. The spectrogram of one unit is shown at the bottom, with the color denoting the power at a given position and spatial frequency. A clear set of peaks in the spectrogram emerges at a fundamental frequency starting at ∼1.1 and at its harmonics. We use a custom algorithm to trace these peaks (see ‘Spectral Decoding’ in Methods) and estimate the gain for each unit. The hippocampal gain, *H*, is estimated as the median spatial frequency across all isolated and mutually coherent units for a given session. (c) Real-time decoding flowchart. Neural data from each tetrode and rat position data from the camera are acquired (see ^28^, for hardware details). Incoming spike times, as detected by Neuralynx spike detection parameters, and positions are added to a temporal buffer. The following operations are performed every 1 s. The temporal buffer is transferred into a spatial queue buffer that accumulates spike times and positions from the previous 6 laps. Velocities are computed from positions, and spikes and positions with velocities < 5 cm/s are eliminated. The remaining spikes and positions are spatially binned (5° width). The spike bins are divided by position bins to create firing rate bins for each tetrode, which are then smoothed and sent to the spectral decoder to estimate H (details in Methods section and Fig. 1 (d-g)). The spectral decoder is able to estimate spatial frequency from the cumulative spatial tuning of all simultaneously recorded cells on a tetrode, extending the success of decoding spatial frequency from cells with such diffuse spatial tuning like interneurons (described in ^6^). The spatial frequency was estimated from each tetrode independently before combining together into 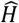, to get independent measurements of the underlying gain and to be robust to noisy estimates on any particular tetrode. (d) Coherence of population gain. *H* was decoded from each unit. If a unit, *i*, were part of a coherent population, its gain *Hi* should equal the population hippocampal gain *H*. For each 6-lap window we computed a coherence error |1*− Hi*/*H*| and computed the average of this value across the session to derive the coherence score for the unit (Data from 1059 units, 66 sessions). Most units have a score very close to zero and very few have values above 0.1 (18 units, range 0.13 – 0.49). (e) Comparison of offline unsorted decoding vs. online sorted decoding in Epoch 2 of the 25 closed-loop sessions. The mean absolute error between these gains remains close to 0, with a few sessions (3/25) showing deviations greater than 0.1.

**Extended Data Fig. 4.**
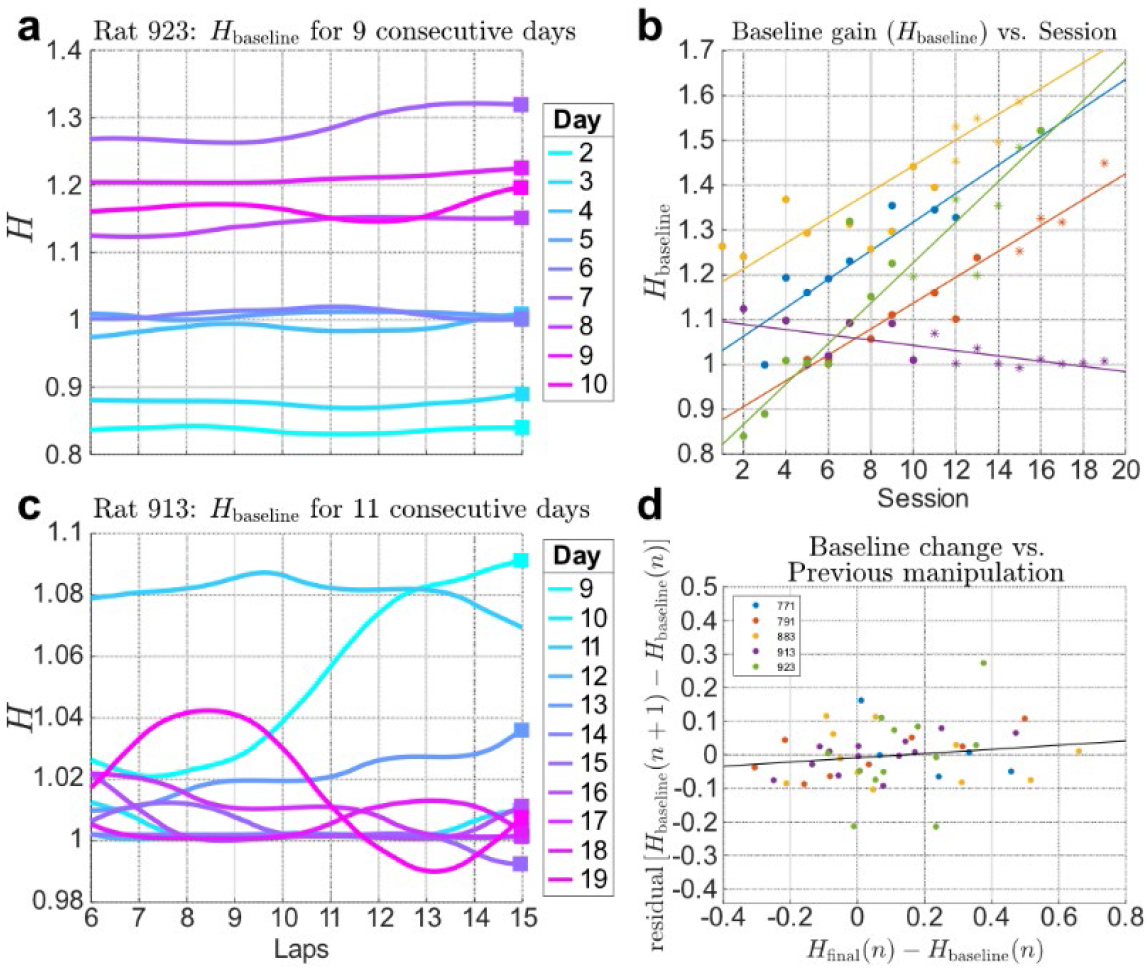
Baseline shift across days. (a) Baseline shift for two rats. The *y* axis is hippocampal gain, *H*, during Epoch 1 (stripes stationary) for 9 consecutive sessions for the same rat. The *x* axis denotes laps on the table. For Rat 923, the value of H at the end of the epoch, *H*_baseline_, steadily increased across sessions but was relatively stable within a session. For Rat 913, *H*_baseline_, decreased over sessions and was less stable within a session. (b) Baseline shift across rats. Data from each rat is plotted in a different color. The *x* axis is the session number of each rat and *y* axis is *H*_baseline_ for that session. Dots denote open-loop sessions and asterisks (*) denote closed-loop sessions. 4/5 rats show a significant positive drift of *H*_baseline_ (Rat 771: slope= 0.032, *r*^2^ = 0.77, *p* = 0.004, *n* = 8; Rat 791: slope= 0.029, *r*^2^ = 0.93, *p* = 5 × 10^−7^, *n* = 12; Rat 883: slope= 0.029, *r*^2^ = 0.73, *p* = 3 × 10^−5^, *n* = 16; Rat 923: slope= 0.045, *r*^2^ = 0.90, *p* = 2 × 10^−7^, *n* = 14) whereas one rat shows a significant negative drift (Rat 913: slope= −0.006, *r*^2^ = 0.45, *p* = 0.005; *n* = 14). Because of these shifts, we subtracted *H*_baseline_ from the dependent measures in our analyses of Figs. 2g, 3h, and 4c. (c) Effect of previous day’s manipulation on baseline shift. We analyzed the effect of the previous day’s manipulation on each session. For each pair of consecutive sessions *n* and *n* + 1, we plotted the gain change induced by stripes on day *n, H*_final_(*n*) *− H*_baseline_(*n*), as the independent variable on the *x* axis. The *y* axis was the change in the baseline to the next day (*n* + 1), minus the linear trend from (b). There was no significant relationship between these variables (*r*^2^ = 0.03, *p* = 0.24, *n* = 49).

**Extended Data Fig. 5.**
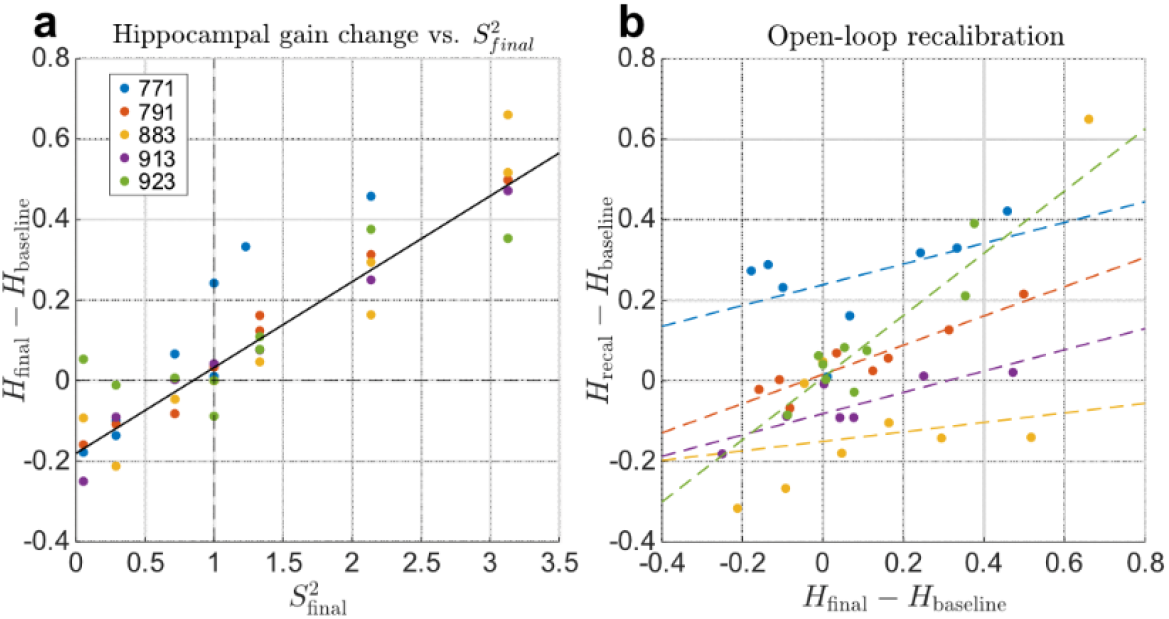
Open-loop control and recalibration. (a) Validity of quadratic fit. Fig. 2g showed the change in hippocampal gain against the final stripe gain and the fits to the relationship using a power law. Since the exponent parameters are close to 2 (individual rat exponents: 1.56, 1.94, 3.31, 1.53, 3.20), here we show a linear fit of hippocampal gain change against 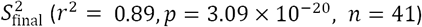. (b) Open-loop recalibration. Although the open-loop manipulation did not result in an *H* value that was stable over a number of laps, the manipulation did result in a modest recalibration of the hippocampal gain after the stripes were extinguished. Similar to Fig. 4d, the x axis shows *H*_final_ and the y axis shows *H*_recal_ (with *H*_baseline_ subtracted from both). For 3/5 rats, the linear relationship is positive and significant (Rat 791, slope= 0.36, *r*^2^ = 0.83, *p* = 0.002; Rat 913, slope= 0.26, *r*^2^ = 0.66, *p* = 0.026; Rat 923, slope= 0.77, *r*^2^ = 0.76, *p* = 0.002). For the other 2 rats, there was a positive, but statistically nonsignificant, relationship (Rat 771, slope= 0.26, *r*^2^ = 0.24, *p* = 0.23; Rat 883, slope= 0.12, *r*^2^ = 0.25, *p* = 0.59).

**Extended Data Fig. 6.**
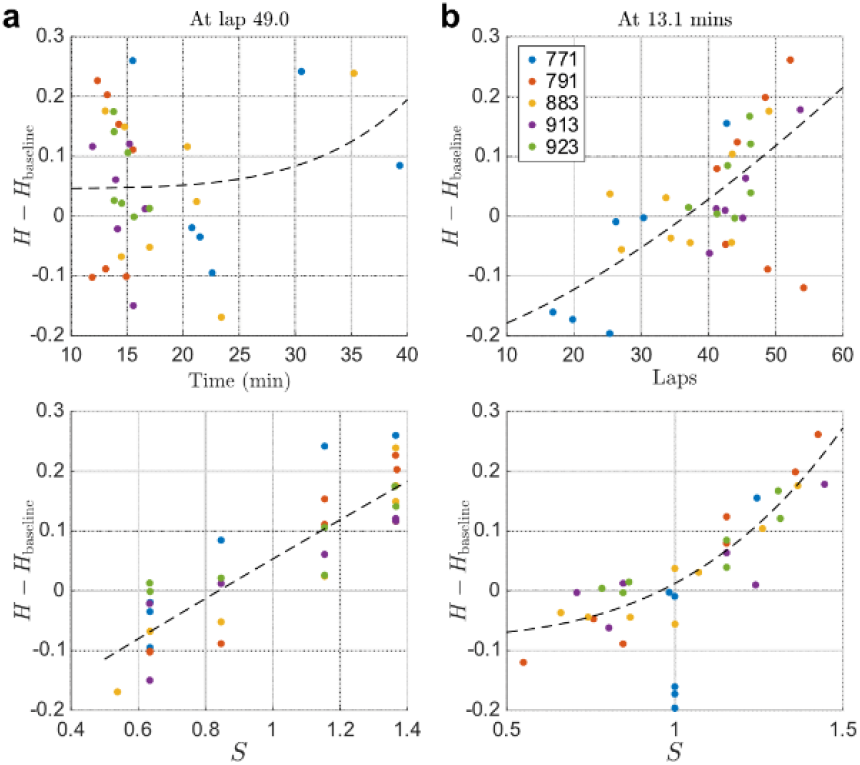
Effect of manipulation time and distance run on hippocampal gain. Because we ramped *S* by a constant rate across experiments, there was a correlation between *S*_final_ and time and distance travelled in our experiments, introducing potential confounding variables. We thus examined the influences of distance run and time spent under stripe manipulation to the change in hippocampal gain, *H* − *H*_baseline_, compared to our stripe gain manipulation *S*. These analyses were run for all open-loop sessions where *S* ≠ 1. (a) For these sessions, we computed the minimum distance rats ran in Epoch 1 and 2 (49 laps, the distance run in the *S*_final_ = 1 ± 2/13 sessions) to equate distance across gain manipulations. At 49 laps after start of Epoch 1, the change in hippocampal gain *H* − *H*_baseline_ is plotted as a function of both time (top) and stripe gain *S* (bottom) for all sessions. Each data point is from a session and colors denote different rats. We fit a power-law curve to both plots (*H* − *H*_baseline_ = *a* + *bx*^*m*^). There is no obvious relationship between *H* − *H*_baseline_ and time (adjusted *r*^2^ = −0.26, *df* = 31), whereas there is a power-law relationship to *S* (adjusted *r*^2^ = 0.72, *df* = 31) similar to Fig. 2(g). (b) For these sessions, we computed the minimum time rats ran in Epochs 1 and 2. These values are variable; however, the minimum value across all sessions was 13.1 minutes, with the 5 rats ranging from 13.1 mins to 19.9 mins. At 17 mins after the start of Epoch 1, the change in hippocampal gain *H* − *H*_baseline_ is plotted as a function of both distance run (top) and stripe gain S (bottom). There is no strong relationship between *H* − *H*_baseline_ and distance (adjusted *r*^2^ = 0.35, *df* = 31) whereas there is a clear power-law relationship to *S* (adjusted *r*^2^ = 0.78, *df* = 31), similar to Fig. 2(g). From these plots, we conclude that the change in H in these experiments is related to S and not to the correlated variables, time and distance travelled.

**Extended Data Fig. 7.**
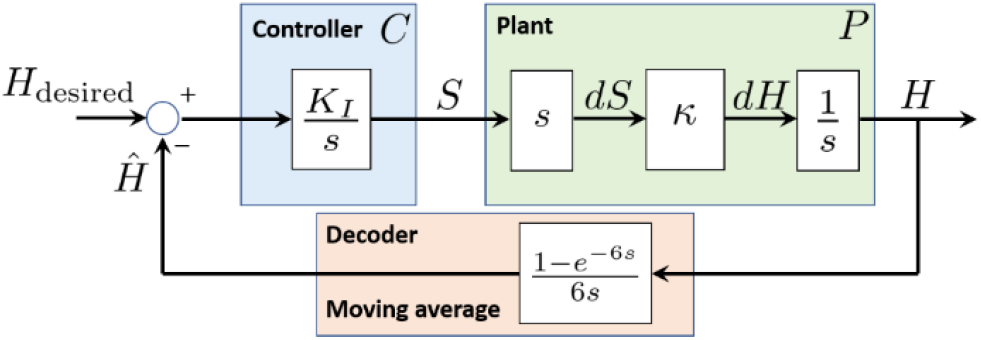
Layout of the system in the Laplace domain. Here, *s* is the Laplace complex frequency variable. Multiplication by *s* denotes differentiation, whereas dividing by *s* denotes integration. *C* is the implementation of our neurally closed-loop controller, namely the transformation from the error 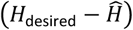 to the stripe gain *S*. In control theoretic terminology, the controlled system (hippocampal circuit) is the “plant” *P*, which transforms the stripe gain *S* into the output *H*. The term 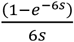 in the feedback loop is the transfer function of a 6-lap moving average, capturing the lag introduced by our online gain decoder. The transfer function of the controller is 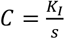 and that of the plant reduces to a constant gain, *P* = *k*.

